# Electrophysiological approaches to understanding brain-muscle interactions during gait: a systematic review

**DOI:** 10.1101/2024.02.27.582247

**Authors:** Maura Seynaeve, Dante Mantini, Toon T. de Beukelaar

**Affiliations:** Movement Control and Neuroplasticity Research Group, Department of Movement Sciences, KU Leuven, 3001 Leuven, Belgium

**Keywords:** Gait – electroencephalography – electromyography – neuromuscular connectivity – corticomuscular coherence – Mobile brain/body imaging

## Abstract

**Objective:** This study systematically reviews the role of the cortex in gait control by analyzing connectivity between electroencephalography (EEG) and electromyography (EMG) signals, i.e. neuromuscular connectivity (NMC) during walking. We aim to answer the following questions: (i) Is there significant NMC during gait in a healthy population? (ii) Is NMC modulated by gait task specifications (e.g. speed, surface, additional task demands)? (iii) Is NMC altered in the elderly or a population affected by a neuromuscular or neurologic disorder?

**Methods:** Following PRISMA guidelines, a systematic search of seven scientific databases was conducted until September 2023.

**Results:** Out of 1308 identified papers, 27 studies met the eligibility criteria. Despite large variability in methodology, significant NMC was detected in most of the studies. NMC was able to discriminate between a healthy population and a population affected by a neuromuscular or neurologic disorder. Tasks requiring higher sensorimotor control resulted in an elevated level of NMC.

**Conclusions:** While NMC holds promise as a metric for advancing our comprehension of brain-muscle interactions during gait, aligning methodologies across studies is imperative.

**Significance:** Analysis of NMC provides valuable insights for the understanding of neural control of movement, development of gait retraining programs and contributes to advancements in neurotechnology.

## 1. Introduction

It has been quantified that we take on average around 6500 steps per day (Paluch et al., 2022). Many of our daily steps occur automatically, often without conscious thought. When we walk, we seamlessly translate the intention to move into a coordinated pattern of activating and deactivating multiple muscle groups. The coordination of these movements is controlled by the central nervous system (CNS). However, the exact mechanisms by which the CNS accomplishes this, are not yet fully understood.

Central pattern generators at the spinal level predominantly control the rhythmic movements of the limb during gait in quadrupedal animals (Brown, 1914; Guertin, 2009). In humans, this control at the spinal level is modulated by supraspinal factors (Takakusaki, 2017). Attempts to understand the cortex’s role in gait target different brain imaging and neuromodulation techniques. For instance, functional magnetic resonance imaging (fMRI) studies found activation in the supplementary motor area, bilateral precentral gyrus, left dorsal premotor cortex, and cingulate motor area during motor imagery of gait (Allali et al., 2014; Wang et al., 2008). Functional infrared spectroscopy (fNIRS) demonstrated an increase in local hemoglobin oxygenation over the sensorimotor cortices and supplementary motor areas during gait, hereby reflecting increased activation in these regions (Kurz et al., 2012; Miyai et al., 2001; Suzuki et al., 2008, 2004). Moreover, changes in corticospinal excitability were established using transcranial magnetic stimulation (TMS) (Bonnard et al., 2002; Capaday et al., 1999; Petersen et al., 1998; Schubert et al., 1997). Although these findings help understand the cortex’s role in gait, these techniques have intrinsic shortcomings. fMRI is non-portable and therefore limited to stationary tasks, e.g. observing or imaging of gait. Although some studies used TMS during active walking, it remains difficult to maintain a precise coil placement while the participant is moving. Moreover, TMS only delivers indirect evidence of cortical activation (Klomjai et al., 2015; Walsh and Cowey, 2000). fNIRS, on the other hand, is portable and can be used dynamically, however, it is characterized by low spatial and temporal resolution and substantial inter-subject variability (Vitorio et al., 2017). Hence, there is a need for a portable brain imaging technique that provides higher resolution.

To address this need, researchers started using electroencephalography (EEG) to study brain dynamics during movements. EEG measures the electrical activity of the brain. Specifically, it records the fluctuations in electrical potential generated by the collective activity of neurons through electrodes placed on the scalp. Its excellent temporal resolution allows for the real-time examination of brain activity during dynamic tasks like walking (Korivand et al., 2023; Walsh and Cowey, 2000). Recent advancements in EEG processing techniques, such as advanced artifact correction methods, have significantly enhanced the quality of recordings during movement (Zhao et al., 2021). Additionally, the utilization of volume conduction models has contributed to improved spatial resolution (Taberna et al., 2021, 2019). Therefore, EEG is well-suited to study temporal and spatial brain dynamics during gait (Korivand et al., 2023).

EEG revealed distinctive neural synchronization patterns in the motor cortex during gait. An increase in EEG power can be detected in double support phases of gait, whereas a decrease in power occurs during single leg stance and swing phase (Gwin et al., 2011; Severens et al., 2012). These changes in EEG power are the result of the synchronizing or desynchronizing of neuronal populations and are called event-related synchronization (ERS) and event-related desynchronization (ERD) respectively. In fact, by synchronizing the rate and timing of their action potentials, distant neuronal populations can communicate with each other (Fries, 2005). In the motor system, this synchronization serves as a mechanism through which upper motor neurons establish communication with the spinal motor neurons. Therefore, this ERD/ERS occurring during gait likely signifies the communication between the primary motor cortex and the spinal motor neurons of the leg muscles via the corticospinal tract. The neural activity of these spinal motor neurons can be indirectly measured using surface electromyography (EMG). Hereby, analyzing the synchronization between EEG and EMG signals can inform us about the way they interact or communicate with each other. In the context of this review, the term ‘neuromuscular connectivity (NMC)’ will be used to express synchronization analyses between the EEG and EMG signal.

NMC is most frequently quantified using coherence, in this case referred to as corticomuscular coherence (CMC) (Liu et al., 2019). Coherence can be considered a frequency domain version of Pearson’s correlation coefficient. In other words, CMC signifies communication between the cortex and muscles. Several studies have reported a weak but significant CMC in the beta frequency range [13-30 Hz] between EMG signals from hand or foot muscles and EEG signals over contralateral sensorimotor regions during sustained isometric muscle contractions (Conway et al., 1995; Gross et al., 2000; Halliday et al., 1998; Salenius et al., 1997). With increasing contraction intensities, CMC tends to increase (Chakarov et al., 2009; Witte et al., 2007), while during movement CMC typically decreases (Baker et al., 1997; Kilner et al., 2000). Despite this movement-related CMC reduction, several studies have identified significant synchronization between the cortex and leg muscles during specific phases of the gait cycle. Therefore, this analysis may increase our understanding of how the CNS controls the coordination of muscles during gait.

Several studies have attempted to quantify the synchronization between EEG and EMG during gait. However, a proper overview of the existing body of literature is currently lacking and therefore hinders definitive conclusions. In this systematic review, we will therefore synthesize all available literature quantifying synchronization between the brain and lower limb muscles during steady-state gait and stepping tasks. This review aims to find an answer to the following questions: (i) is there significant NMC during gait in a normal, healthy population? (ii) Is NMC modulated by gait task specifications (e.g. speed, surface, additional task demands)? (iii) Is NMC altered in the elderly or a population affected by a neuromuscular disorder?

## 2. Methods

### 2.1. Search strategy

The search strategy aimed to identify relevant studies investigating neuromuscular connectivity (NMC) during gait. A comprehensive search was conducted across seven electronic databases including PubMed, Embase, Scopus, Web of Science, Cochrane Library, SportDiscus, and IEEE Xplore. The search query utilized a combination of three different concepts that were combined with the AND Boolean operator. The first concept was electromyography, searched for with the keywords ‘electromyogr*’, ‘EMG’, and ‘sEMG’. The second concept was electroencephalography, searched for with the keywords ‘electroencephalogr*’, ‘EEG’, and ‘hdEEG’. The third concept was gait, searched for with the keywords ‘gait’, ‘walk*’, and ‘locomotion’. The search was limited to articles published up until September 2023.

### 2.2. Eligibility criteria

All studies which met the following inclusion criteria were considered for the current systematic review:

- To include the specified query in the abstract and/or title and/or in the keywords
- To involve the simultaneous use of EMG and EEG during a gait or stepping task on a stable, level surface
- To involve quantification of the synchronization between EEG and EMG signal
- To be indexed in at least one of the screened databases
- To be available in English
- To be a full article (i.e., no reviews, editorials or conference abstracts)

### 2.3. Study selection

After removing duplicates, titles and abstracts were independently screened by two reviewers. Next, retrieved studies were separately evaluated by the reviewers in accordance with the eligibility criteria. Any disparities between the authors were discussed with a third reviewer until a consensus was reached.

### 2.4. Data extraction and synthesis

Of the studies that were selected based on the eligibility criteria, the following information was extracted: authors, publication date, study aim, participant characteristics (sample size, age, and gender), gait task characteristics (speed, surface, duration, additional demands), EMG-set and muscles measured, EEG set-up, EMG and EEG preprocessing steps, synchronization measure and the gait phases examined. In case of discrepancies between reviewers during data extraction, these discrepancies were thoroughly discussed with a third reviewer until consensus was reached. The results are synthesized in table 1.

**Table 1.** Study characteristics.

|  | ***Participants***<br>***(sample size,***<br>***mean age)*** | ***Gait task*** | ***EMG set-up and***<br>***preprocessing*** | ***EEG set-up and***<br>***preprocessing*** | ***EMG-EEG***<br>***synchronization***<br>***measure, frequency***<br>***ranges of interest***<br>***and gait phase of***<br>***interest*** |
| --- | --- | --- | --- | --- | --- |
| **Petersen et al., 2012** | Healthy young (9, F: 5, age: 23.4 ± 4.1) | Treadmill walking at 3.5-4 km/h and 1 km/h | Unilateral TA<br>Freq. 1-1000 Hz<br>Rectified signal | 28-channel montage<br>Freq. 1-250 Hz<br>ICA artifact removal<br>Sensor-level analysis (Cz) | Coherence<br>[8-13 Hz] [24-40 Hz]<br>800ms before HS – 200ms after HS |
| **de Tommaso et al., 2015** | Healthy (17, F: 12, age range: 18-65) | Overground walking and cognitive dual-task walking at preferred speed | Bilateral TA, GCL,<br>Freq. 10-90 Hz<br>Unrectified signal | 21-channel montage<br>Freq. 7-30 Hz<br>ICA artifact removal<br>Sensor-level analysis | Coherence<br>[8-13 Hz] [13-30 Hz] |
| **Winslow and Brantley, 2016** | Healthy young (1, F: 0, age: 31) | Overground walking at preferred speed | Unilateral TA<br>Freq. 1-450 Hz<br>Unrectified signal | 64-channel montage<br>Freq. 0.1-500 Hz<br>ICA artifact removal<br>Sensor-level analysis (Cz) | Coherence<br>[20-40 Hz]<br>Complete gait cycle |
| **Brantley and Winslow, 2016** | Healthy young (1, F: 0, age: 31) | Overground walking at preferred speed | Bilateral TA, GCM, RF, VL, ST<br>Freq. 30-50 Hz<br>Unrectified signal | 64-channel montage<br>Freq. 0.1-6 Hz<br>ASR artifact removal<br>Sensor-level analysis (Cz, C1, C2) | Coherence<br>[1-6 Hz]<br>Complete gait cycle |
| **Storzer et al., 2016** | Healthy young (15, F: 6, age: 24.9 ± 3) | Overground walking at 40 bpm at preferred speed | Bilateral TA, RF, BF<br>Freq. 20-450 Hz<br>Rectified signal | 18-channel montage<br>Freq. 1-100 Hz<br>ICA artifact removal<br>Sensor-level analysis (Cz) | Power correlation<br>[8-13 Hz] [24-40 Hz]<br>Complete gait cycle |
| **Artoni et al., 2017** | Healthy young (11, age: 30 ± 4) | Treadmill walking at 3.5 km/h | Bilateral TA, VM, BF<br>Freq. 2-500 Hz<br>Unrectified signal | 64-channel montage<br>Freq. 0-256 Hz<br>ICA + ASR artifact removal<br>Source-level analysis | Phase coupling<br>[1-45 Hz]<br>HS excluded |
| **Roeder et al., 2018** | Healthy young (24, F: 12, age: 25.9 ± 3.2) | Overground walking and treadmill walking at preferred speed (4.2 ± 0.4km/h), barefoot | Bilateral TA<br>Freq. 0-10 Hz<br>Rectified signal | 10-channel montage<br>Freq. 1-500 Hz<br>ICA artifact removal<br>Sensor-level analysis (C3-F3, C4-F4) | Coherence<br>[4-8 Hz] [8-13 Hz] [13-30 Hz] [30-50 Hz]<br>800ms before HS – 200ms after HS |
| **Jensen et al., 2018** | Healthy young (16, F: 10, age: 23 ± 5) | Treadmill walking and visually guided walking at 2.2 km/h | Unilateral TA, SOL, GCM<br>Rectified signal | 2-channel montage<br>Freq. 0.5-70 Hz<br>No artifact removal<br>Sensor-level analysis (Cz) | Coherence<br>[8-13 Hz] [13-30 Hz] [30-70 Hz]<br>Swing phase (TA) and stance phase (SOL/GCM), HS excluded |
| **Jensen et al., 2019a (main study)** | Healthy young (11, F: 6, age: 24.9 ± 2.8) | Treadmill walking at 3.6 km/h | SOL, GCM<br>Rectified signal | 2-channel montage<br>Freq. 5-500 Hz<br>No artifact removal<br>Sensor-level analysis (Cz) | Coherence<br>[5-50 Hz]<br>HS excluded |
| **Jensen et al., 2019b (control experiment)** | Healthy young (10, F: 6, age: 26.3 ± 4.5) | Treadmill walking at 3.6 km/h | SOL, GCM<br>Unrectified signal | 64-channel montage<br>Freq. 0-1024 Hz<br>No artifact removal<br>Sensor-level analysis | Coherence<br>[30-64 Hz]<br>HS excluded |
| **Günther et al., 2019** | Healthy old (3, F: 1, age: 65.7 ± 14.2)<br>PD-FOG (5, F: 2, age: 68.8 ± 9)<br>PD+FOG (4, F: 0, age: 64.3 ± 8.2) | Overground figure-eight walking | Bilateral TA, GCM<br>Freq. 0-10 Hz<br>Unrectified signal | 32-channel montage<br>Freq. 0.1-128 Hz<br>ICA artifact removal<br>Sensor-level analysis (C3-C4) | Power correlation<br>[4-8 Hz] [8-13 Hz] [13-30 Hz] |
| **Li et al., 2019** | Healthy young (30, F: 0, age: 24 ± 2.32) | Overground walking and walking with exoskeleton at preferred speed | Unilateral TA, GCL, RF, ST<br>Freq. 2-125 Hz<br>Unrectified signal | 62-channel montage<br>Freq. 0.5-45 Hz<br>ICA artifact removal<br>Sensor-level analysis | Coherence<br>[2-4 Hz] [4-8 Hz] [8-13 Hz] [13-30 Hz] [30-50 Hz] |
| **Spedden et al., 2019** | Healthy young (15, F: 8, age: 22.1 ± 1.7)<br>Healthy old (15, F: 8, age: 68.3 ± 2.7) | Treadmill walking and visually guided walking. Step length and speed normalized to leg length (healthy young: 2.13 ± | Unilateral TA<br>Freq. 4-80 Hz<br>Rectified signal | 64-channel montage<br>Freq. 4-80 Hz<br>ICA artifact removal<br>Sensor-level analysis (Cz) | Coherence<br>[13-30 Hz] [30-50 Hz]<br>Swing phase, 650 – 50ms before HS, HS excluded |
| | | 0.12 and old: $2.15 \pm 0.16$ km/h) | | | |
| **Hoxha et al., 2019** | Healthy (8, F: 6, age: $51 \pm 6$)<br>MS (8, F: 6, age: $53 \pm 6$) | Treadmill walking at preferred speed (healthy: $2.156 \pm 0.3$ km/h and MS: $1.787 \pm 1.1$ km/h) | Bilateral TA, SOL, GC | 64-channel montage<br>Freq. 3-50 Hz<br>ICA artifact removal<br>Source-level analysis | Phase coupling [13-30 Hz]<br>Complete gait cycle |
| **Chen et al., 2019** | Stroke control group (9, F: 1, $50.33 \pm 9.77$)<br>Stroke experimental group (9, F: 0, $54.67 \pm 8.32$) | Treadmill walking at preferred speed (range: 2.08 – 2.56 km/h) | Unilateral TA | 32-channel montage<br>Freq. 1-50 Hz<br>ICA artifact removal<br>Sensor-level analysis (all channels) | Coherence [8-13 Hz] [30-50 Hz]<br>Swing phase |
| **Short et al., 2020** | Healthy adolescents (12, F: 8, $14.8 \pm 3$)<br>CP (9, F: 7, $16 \pm 2.7$) | Treadmill walking at preferred speed (3.2 – 3.6 km/h) | Bilateral TA, SOL, GCM, RF, VL, ST, PL, HL<br>Freq. 5-35 Hz<br>Rectified signal | 64-channel montage<br>Freq. 1-500 Hz<br>ICA + ASR artifact removal<br>Source-level analysis | Coherence [2-4 Hz] [4-8 Hz] [8-13 Hz] [13-30 Hz] [30-50 Hz]<br>Complete gait cycle |
| **Roeder et al., 2020** | Healthy young (24, F: 12, age: $25.9 \pm 3.2$)<br>Healthy old (24, F: 12, age: $65.1 \pm 7.8$)<br>PD (21, F: 8, age: $67.4 \pm 7.3$) | Overground and treadmill walking at preferred speed (healthy young: $4.16 \pm 0.12$, healthy old: $4.00 \pm 0.11$, PD: $3.97 \pm 0.12$), barefoot | Bilateral TA<br>Freq. 20-45 Hz<br>Rectified signal | 10-channel montage<br>Freq. 1-500 Hz<br>ICA artifact removal<br>Sensor-level analysis (C3-F3, C4-F4) | Coherence [4-8 Hz] [8-13 Hz] [13-30 Hz] [30-50 Hz]<br>800ms before HS – 200ms after HS |
| **Yokoyama et al., 2020** | Healthy young (15, F: 0, age: $26.7 \pm 7.5$)<br>Healthy old (9, F: 0, age: $64.9 \pm 6.3$)<br>PD: (10, F: 0, age: $61.6 \pm 6.3$) | Overground walking at preferred speed ($4.104 \pm 0.504$ km/h for PD, $4.572 \pm 0.504$ km/h for healthy old, and $4.5 \pm 0.468$ km/h for healthy young), barefoot | Bilateral TA, GCM<br>Freq. 0-64 Hz<br>Rectified signal | 20-channel montage<br>Freq. 2-200 Hz<br>ASR artifact removal<br>Sensor-level analysis (Cz) | Phase coupling [8-13 Hz] [13-30 Hz] [30-50 Hz]<br>Swing phase (TA) and stance phase (GCM) |
| **Gennaro et al., 2020a** | Healthy young (9, F: 5, age: $26 \pm 3$)<br>Healthy old (9, F: 3, age: $73 \pm 6$) | Overground figure eight at preferred speed | Bilateral TA<br>Freq. 20-250 Hz<br>Rectified signal | 64-channel montage<br>Freq. 1.5-48 Hz<br>ICA + ASR artifact removal<br>Sensor-level analysis (Cz) | Coherence [13-30 Hz] [30-40 Hz]<br>Swing phase, 650 – 50ms before HS, HS excluded |
| **Gennaro et al., 2020b** | Healthy old (11, F: 6, age: $72 \pm 4$)<br>Sarcopenia (11, F: 9, age: $75 \pm 7$) | Overground figure eight at preferred speed ($3.888 \pm 0.756$ km/h) | Bilateral TA, SOL, GCM, GCL, RF, VL, VM, BF<br>Freq. 0-20 Hz<br>Rectified signal | 64-channel montage<br>Freq. 1.5-48 Hz<br>ICA + ASR artifact removal<br>Sensor-level analysis (Cz) | Coherence [13-30 Hz] [30-48 Hz]<br>Swing phase, 650 – 50ms before HS, HS excluded |
| **Chen et al., 2021** | Healthy young (6, F: 3, age range: 24-26) | Overground slow walking at 60 bpm and fast walking at 120 bpm | Bilateral TA, VM, ST<br>Freq. 0.5-50 Hz<br>Unrectified signal | 64-channel montage<br>Freq. 0.5-50 Hz<br>ICA artifact removal<br>Sensor-level analysis | Mutual information [4-8 Hz] [8-13 Hz] [13-30 Hz] [30-50 Hz] |
| **Wei et al., 2021** | Healthy young (9, F: 3, age range: 23-26) | Treadmill slow walking at 1.4 km/h, normal walking at 2 km/h and fast walking at 2.6 km/h | Bilateral TA, VM, BF<br>Freq. 30-450 Hz<br>Unrectified signal | 32-channel montage<br>Freq. 0.5-50 Hz<br>ICA artifact removal<br>Sensor-level analysis | Mutual information [13-30 Hz]<br>Complete gait cycle |
| **Manuel Mayor-Torres et al., 2022** | Healthy young (3, F: 2, age: $36 \pm 12.12$)<br>Stroke (3, F: 1, age: $57 \pm 8.71$) | Overground walking and exoskeleton walking | Bilateral TA, SOL, RF, ST<br>Freq. 5-100 Hz<br>Unrectified signal | 32-channel montage<br>Freq. 0.1-100 Hz<br>ICA + ASR artifact removal<br>Sensor-level analysis | Coherence<br>Complete gait cycle |
| **Caffi et al., 2022** | Healthy young (16, F: 9, age IQR: 24 – 25.25)<br>Healthy old (13, F: 4, age IQR: 68 – 71) | Overground walking, cognitive dual task walking and targeted walking at preferred speed | Bilateral TA, SOL, RF, SM<br>Freq. 0-120 Hz<br>Rectified signal | 8-channel montage<br>Freq. 1-120 Hz<br>ASR artifact removal<br>Sensor-level analysis (C3, C4) | Coherence [15-30 Hz] [30-60 Hz]<br>HS excluded, only single support phases |
| **Zhao et al., 2022** | Healthy young (24, F: 14, age range: 22-31) | Treadmill walking at preferred speed | TA, GC, VM, BF<br>Freq. 1-100 Hz<br>Unrectified signal | 128-channel montage<br>Freq. 1-80 Hz<br>ICA artifact removal<br>Source-level analysis | Power correlation [8-13 Hz] [13-30 Hz] [30-50 Hz]<br>Complete gait cycle |
| **Arunganesh et al., 2022** | Healthy young (10, F: 5, age range: (18-31)) | Overground walking at preferred speed | Bilateral TA | 64-channel montage<br>Freq. 0.1-500 Hz<br>Sensor-level analysis (Cz, C1, C2) | Mutual information [0.4-45 Hz] |
| **Roeder et al., 2023** | Healthy young (24, F: 12, age: $25.9 \pm 3.2$)<br>Healthy old (24, F: | Overground walking at preferred speed (3.3-4.8 km/h), barefoot | Bilateral TA, SOL, GCM, GCL<br>Freq. 0-20 Hz<br>Rectified signal | 10-channel montage<br>Freq. 0.5-70 Hz<br>ICA artifact removal<br>Sensor-level analysis | Coherence<br>300ms before HS – 300ms after HS |
| | 12, age: $65.1 \pm 7.8$)<br>PD (21, F: 8, age:<br>$67.4 \pm 7.3$) | | | (C3-F3, C4-F4) | |
| **Spedden et al., 2023** | Healthy young (31, F: 9, age: $26 \pm 4$) | Overground visually guided steps and self guided steps | TA<br>Freq. 0-5 Hz<br>Rectified signal | 64-channel montage<br>Freq. 1-256 Hz<br>ICA artifact removal<br>Source-level analysis | Coherence<br>[5-15 Hz] [25-45 Hz]<br>Swing phase of single step |

## 3. Results

### 3.1. Selected papers

A total of 1308 papers were identified through the screening process across multiple databases. Among these, 618 papers were identified as duplicates and 690 studies progressed to the screening phase. After an evaluation of the title and abstract of these studies against the pre-defined eligibility criteria, 29 papers were deemed eligible and included in the review. Following the assessment of the full articles, two additional studies were excluded as the walking task was not performed on a flat surface. Instead, the task involved climbing stairs or walking on a balance beam. The findings of our research are presented in the following sections. A visual representation of the screening and inclusion process can be found in figure 1, which illustrates the steps outlined in the PRISMA flow chart.

**Figure 1.**
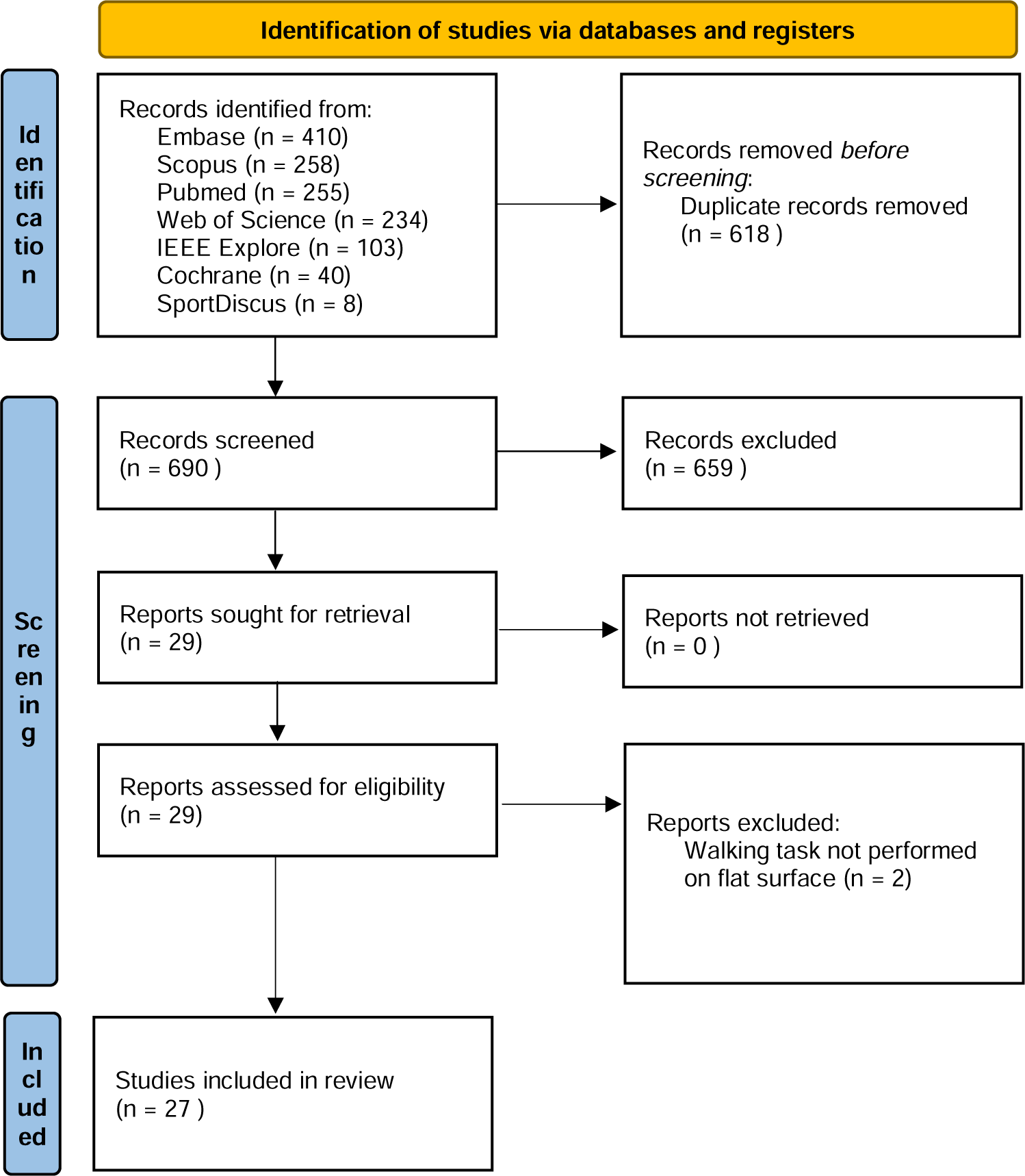
Preferred items for systematic reviews and meta-analyses (PRISMA) flow diagram.

### 3.2. Study objectives

The different objectives of the included studies are visualized in figure 2. The earliest studies quantifying NMC had the main objective of improving our understanding of the neural control of gait. Subsequently, the development of the neurotechnology field led to an increased interest in NMC analysis, seeking to leverage it for the enhancement of neural machine interfaces and neuroprosthetic systems. Most studies, however, focused on comparing either population groups or conditions, with three studies looking into the interaction effects of these two factors. A small number of studies tested a new NMC analysis technique or tested the reliability of an existing technique. Finally, there was one study that used NMC to examine gait performance change after a treadmill training program. In sum, this broad range of study objectives originating from various fields illustrates the growing and heterogeneous interest in NMC analysis.

**Figure 2.**
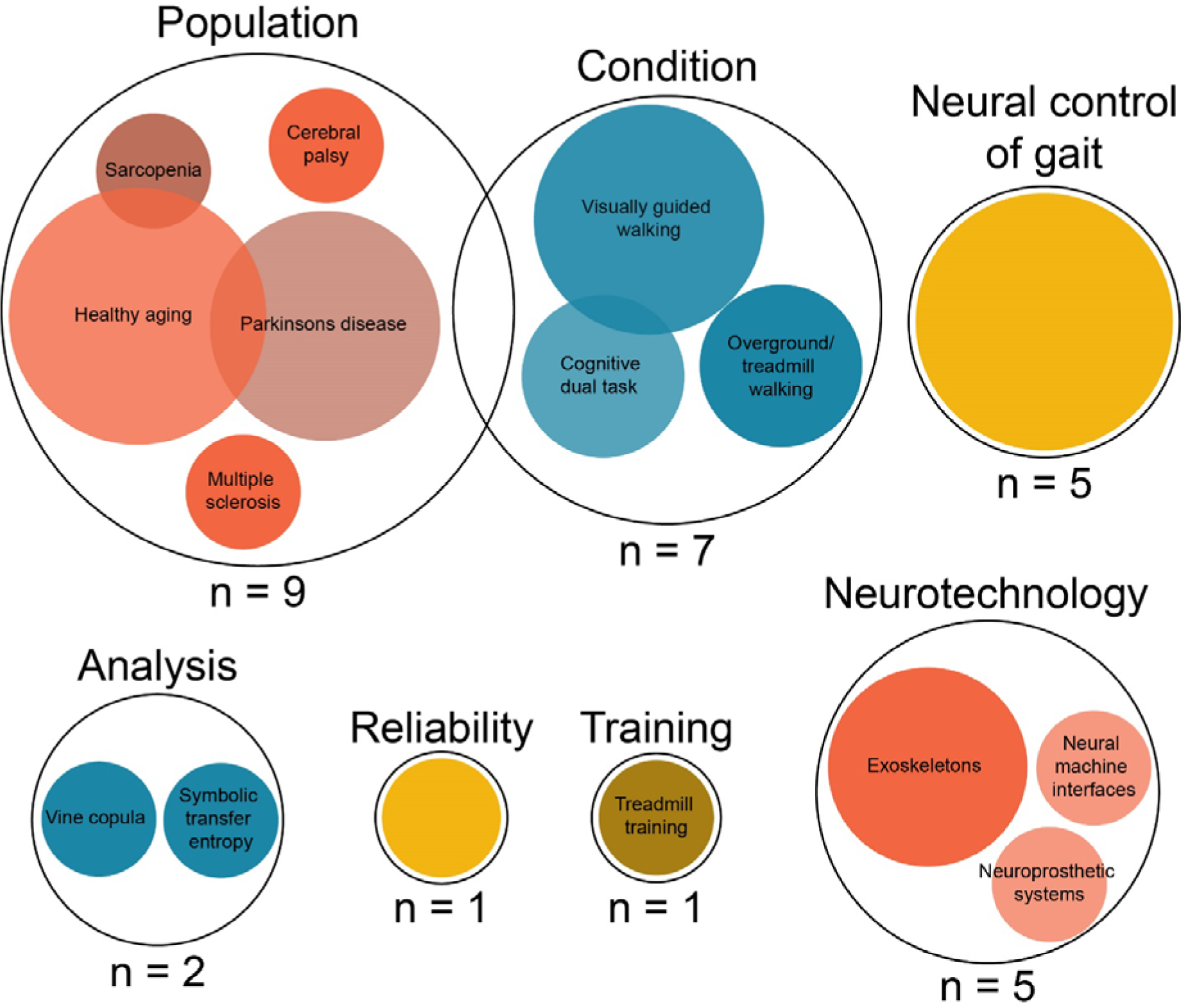
Study objectives. Overlapping circles represent studies having different research objectives.

### 3.3. Participants

The included studies encompass various population groups. Figure 3 offers an overview of the subjects’ age range and sample sizes. Additionally, it distinguishes between healthy participants and those with a neurological or neuromuscular disorder. When studies provided only age range data, the graph represents the average of the lower and upper boundaries of this range.

**Figure 3.**
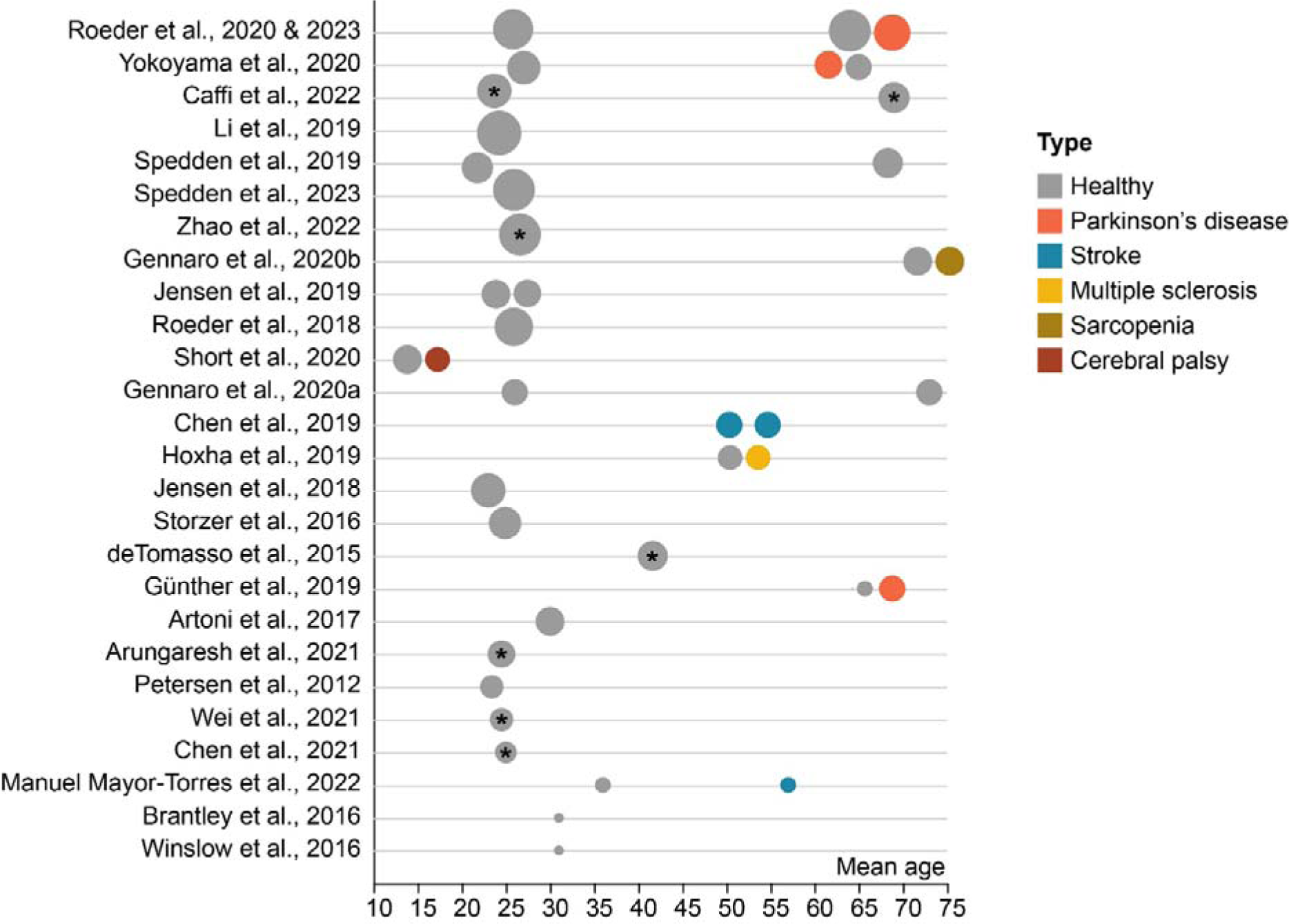
Sample sizes and age distribution across participants. The size of the dots resembles the group sample size. Studies are sorted based on the total sample size. *For the given samples, only age ranges were reported. In this context, the mean of each age range, calculated as the average of the lower and upper bounds, was selected for visualization in the figure.

The age of the subjects ranged from 15 to 75 years old. Particular emphasis was given to two distinct age groups: young adults (aged 22-31 years) and older adults (aged 61-75 years). There is limited representation of middle-aged adults (35 – 60 years), comprising only four studies. Among the 28 studies analyzed, gender information is absent in one study. 45% of participants across the remaining 26 studies are female.

Comparing NMC between young and older individuals during gait yielded conflicting results. Specifically, two studies reported that in the beta and gamma frequency ranges, NMC was higher in younger individuals during both the early swing and double support phases. However, in contrast, three other studies found no significant differences in NMC between the two age groups. Furthermore, two of the studies identified a significant interaction effect between age group and task (Caffi et al., 2022; Spedden et al., 2019). Specifically, when a dual task was introduced alongside regular walking, older individuals exhibited an increase in NMC, whereas younger individuals did not show the same pattern.

Apart from the aging population, studies also investigated participants with neurological or neuromuscular diseases. Nine studies included individuals with various neuromuscular disorders. Of these, four studies included Parkinson’s disease (PD) patients, two studies stroke survivors, one study participants with cerebral palsy (CP), one study multiple sclerosis (MS) and one study sarcopenia. The two most recent studies of Roeder et al. used the same sample of participants and are therefore only represented once in the graph (Roeder et al., 2023, 2020). Out of 27 studies, 12 studies reported to have excluded one or more participants due to various reasons including extensive artifacts in EEG signal or problems associated with the footswitch recordings.

When comparing PD patients to age-matched healthy controls during continuous walking in terms of NMC, conflicting results have emerged. One group observed a significant reduction in NMC among PD patients (Yokoyama et al., 2020). Two other studies, both conducted on the same dataset, found no discernible differences between PD patients and healthy elderly individuals (Roeder et al., 2023, 2020).

One study revealed reduced and more local beta NMC in stroke patients compared to healthy controls during overground walking (Manuel Mayor-Torres et al., n.d.). Similarly, lower NMC was observed in individuals with MS when compared to healthy controls (Hoxha et al., 2019). In the case of CP, patients exhibited unique patterns: they displayed delta band NMC in the non-dominant motor cortex, unlike typically developing children. Furthermore, CP patients showed NMC with the hallucis longus muscle on the dominant side, which is predominantly affected in unilateral CP. NMC to this muscle was not observed in typically developing individuals (Short et al., 2020). Additionally, NMC analysis proved effective in distinguishing between sarcopenic and non-sarcopenic older adults, with sarcopenic adults showing a higher NMC value (Gennaro et al., 2020). This highlights the diagnostic potential of NMC in assessing muscle health in aging populations.

In terms of sample size, seven studies have a limited total sample size of 10 participants or less. Within this subset, two studies are case reports. Furthermore, nine studies feature a moderately larger sample size, ranging from 11 to 20 participants. The remaining 11 studies stand out by featuring a more substantial sample size, examining more than 20 participants. In their two most recent studies, Roeder et al. examined the same group of participants. This is the largest group examined in this review and encompasses 22 young and healthy individuals, 24 healthy elderly controls, and 20 patients diagnosed with PD (Roeder et al., 2023, 2020).

### 3.4. Gait tasks

Information about the gait task and the determination of speed can be found in figure 4. Among the 27 studies considered, nine exclusively focused on a single gait condition, while the remaining studies examined multiple walking conditions. Apart from one study that focuses solely on taking steps, the other studies examined continuous steady-state walking. This could be either on a treadmill or overground. Interestingly, Roeder et al. compared NMC between overground and treadmill walking (Roeder et al., 2018). In 55% of the normal gait conditions, participants walked at a self-selected, preferred speed. In the other studies, the speed or cadence was determined for the participant. Walking speed overall ranged from 1 to 4.8 km/hour.

**Figure 4.**
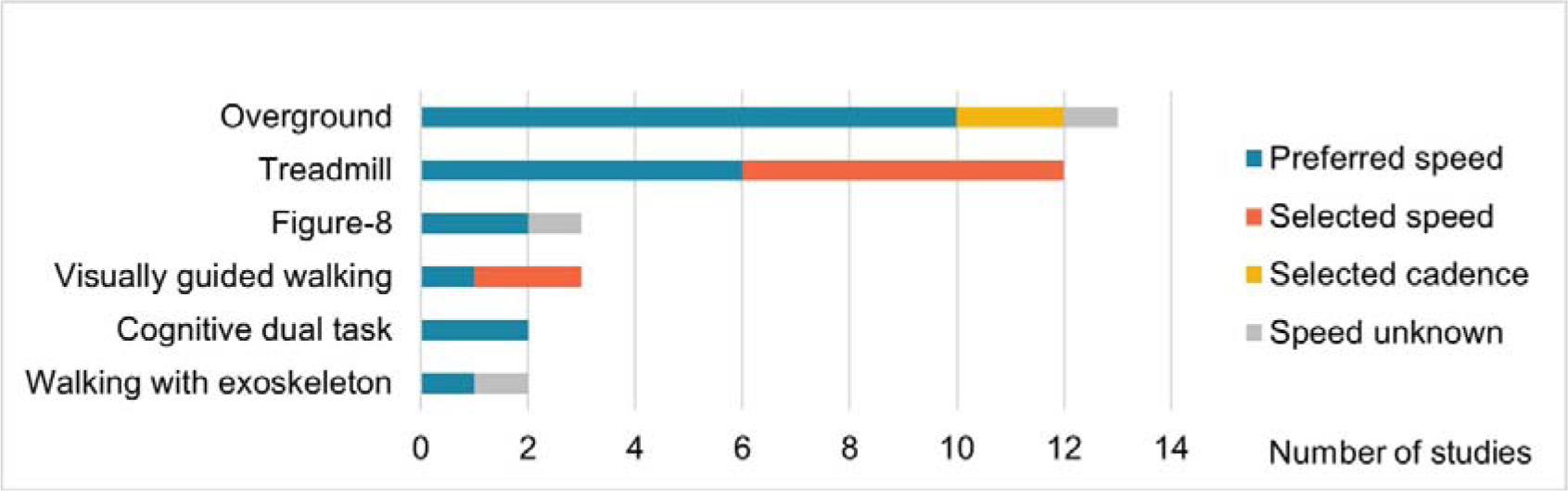
Walking task and speed.

In addition to the predominant steady-state gait, several studies encompassed various other gait tasks, including dual-task walking, visually guided walking, walking backward, walking with intermittent stops, walking in a figure-8 pattern, walking with the assistance of an exoskeleton, and walking on stairs or ramps. In this review, we will only discuss steady-state gait and stepping tasks that are executed on a flat, stable surface.

The different walking conditions offer the opportunity to compare NMC between different gait task demands. Three studies investigated the difference between normal walking and visually guided (VG) walking. During this task, participants had to adjust their step length to hit visual targets either on a wall in front of them or on the floor (Caffi et al., 2022; Jensen et al., 2018; Spedden et al., 2019). In this study, Jensen et al. observed an increase in NMC during VG walking at beta and gamma frequencies, although statistical significance was not reached (Jensen et al., 2018). Subsequent research by Spedden et al. did find a significant increase in NMC during VG walking and this finding was later confirmed by Caffi et al. (Caffi et al., 2022; Spedden et al., 2019).

Two studies conducted by Roeder et al. compared treadmill walking to overground walking. In both cases, they observed that corticomuscular coherence was significantly higher during overground walking compared to treadmill walking. Additionally, three studies independently found distinct patterns related to walking speed. They discovered that walking slowly resulted in higher NMC compared to walking at a faster pace (Chen et al., 2021; Petersen et al., 2012; Wei et al., 2021). Meanwhile, Chen et al. reported that walking forward exhibited higher NMC when contrasted with walking backward (Chen et al., 2021).

Two studies investigated the impact of a cognitive dual task on NMC. In the study by de Tommaso et al. NMC decreased when a dual task was performed (De Tommaso et al., 2015). Conversely, in the study of Caffi et al. an increase in NMC was observed (Caffi et al., 2022). Intriguingly, both studies identified an age-related component, with the modulation being more pronounced in elderly individuals.

### 3.5. EEG set-up and processing

We analyzed the set-up and processing techniques for the EEG signal acquisition (Figure 5). Two studies used a limited number of two electrodes, and Zhao et al. opted for a set of 128 electrodes (Zhao et al., 2022). Two different methods for EEG artifact removal were primarily used. First, studies used Independent Component Analysis (ICA), a signal processing technique used for separating a multivariate signal (e.g., EEG data) into additive, independent components. In the context of EEG artifact removal, ICA is applied to identify and isolate independent components that correspond to artifacts (e.g., eye blinks, muscle activity) from those that represent the underlying neural signals. By separating the signal into independent components, researchers can remove or correct the artifact components, thereby improving the quality and interpretability of the EEG data. The second used technique for artifact removal is Artifact Subspace Reconstruction (ASR). ASR is an advanced method for automatically detecting and removing artifacts in EEG data. ASR works by modeling the artifact subspace using a statistical approach, allowing it to recognize and remove various types of artifacts, including abrupt and non-stationary ones. ASR can be particularly effective for cleaning EEG data by identifying and estimating the artifact components without prior knowledge of the artifact type. It provides a data-driven and automated approach to enhancing the quality of EEG recordings. Twenty-one out of 27 studies included in this review used ICA, while eight used ASR. Of these, five studies used a combination of ASR and ICA, while four studies reported no artifact correction methods.

**Figure 5.**
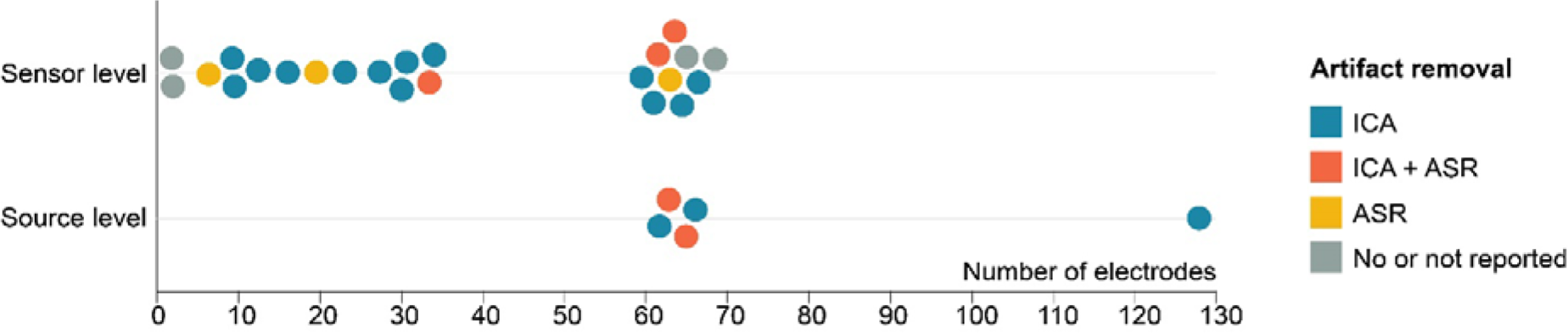
EEG set-up, artifact removal techniques and type of analysis per study. ICA, independent component analysis; ASR, Artifact subspace reconstruction.

Five studies conducted EEG data analysis in the source space, which involves mapping the recorded electrical signals to specific brain regions responsible for generating them. In the study of Zhao et al. an individual MRI of the participant was used to enhance the precision of source localization (Zhao et al., 2022).

Various approaches have been employed for the selection of electrodes of interest in the reviewed studies. Most of these studies utilized the central Cz electrode as a reference point to calculate NMC. Other studies concentrated on electrodes situated on either side of the brain. Interestingly, some investigations examined the spatial distribution of NMC across the entire brain. Two of these clearly reported NMC to be the largest over the Cz electrode (Jensen et al., 2019; Petersen et al., 2012).

### 3.6. EMG set-up and processing

The different muscles examined in the studies can be found in figure 6. Apart from one study, the tibialis anterior muscle (TA) activity was measured in all studies exploring brain-muscle connectivity during gait. In addition, a significant emphasis was placed on the triceps surae complex. Eight studies focused on the soleus muscle (SOL), while thirteen studies investigated one or both heads of the gastrocnemius muscle (GC). The peroneus longus (PL) and the hallucis longus (HL) were each examined in one study.

**Figure 6.**
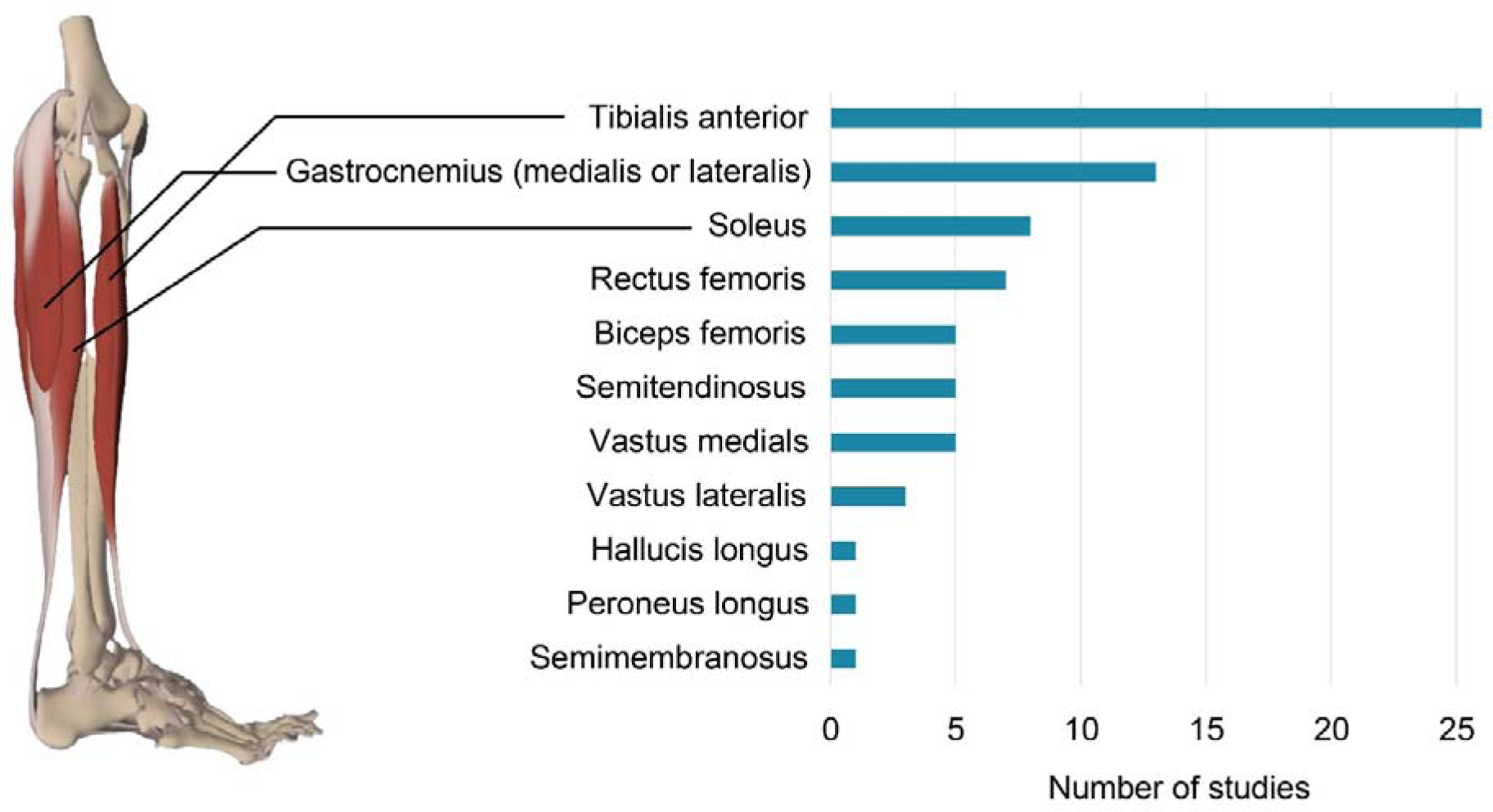
Number of studies conducted for each muscle. © Pharma Intelligence UK Ltd (trading as Primal Pictures), 2023. www.primalpictures.com www.anatomy.tv

Exploration of muscles in the proximal portion of the lower leg was comparatively less common. Specifically, seven studies focused on the rectus femoris muscle. Seven studies investigated the vasti, of which two focused on the vastus lateralis, four on the vastus medialis, and one on both components. Eleven papers examined the hamstring muscles, of which five focused on the semitendinosus, another five on the biceps femoris, and one on the semimembranosus.

Considering EMG analysis, there has been some controversy regarding the rectification of the EMG signal. Fourteen studies applied rectification to the EMG signal, while 11 studies did not.

### 3.7. NMC methods

Across studies, different methods have been adopted to calculate NMC (Figure 7). Corticomuscular coherence, a measure that quantifies the frequency-domain relationship between these two physiological signals, is the most frequently used approach. To calculate coherence, these signals are broken down into distinct frequency components through the Fourier Transform. The cross-spectral density between EEG and EMG is computed to gauge the strength of their relationship at specific frequencies. Concurrently, power spectral density for EEG and EMG separately is determined to ascertain the signal strength in each frequency band. Coherence is then calculated by dividing the cross-spectral density with the product of the power spectral densities of EEG and EMG signals, respectively. This provides a numerical value between zero and one for each relevant frequency range. Higher coherence values are thought to indicate robust synchronization between brain and muscle activities, while lower values on the contrary imply weaker synchronization.

**Figure 7.**
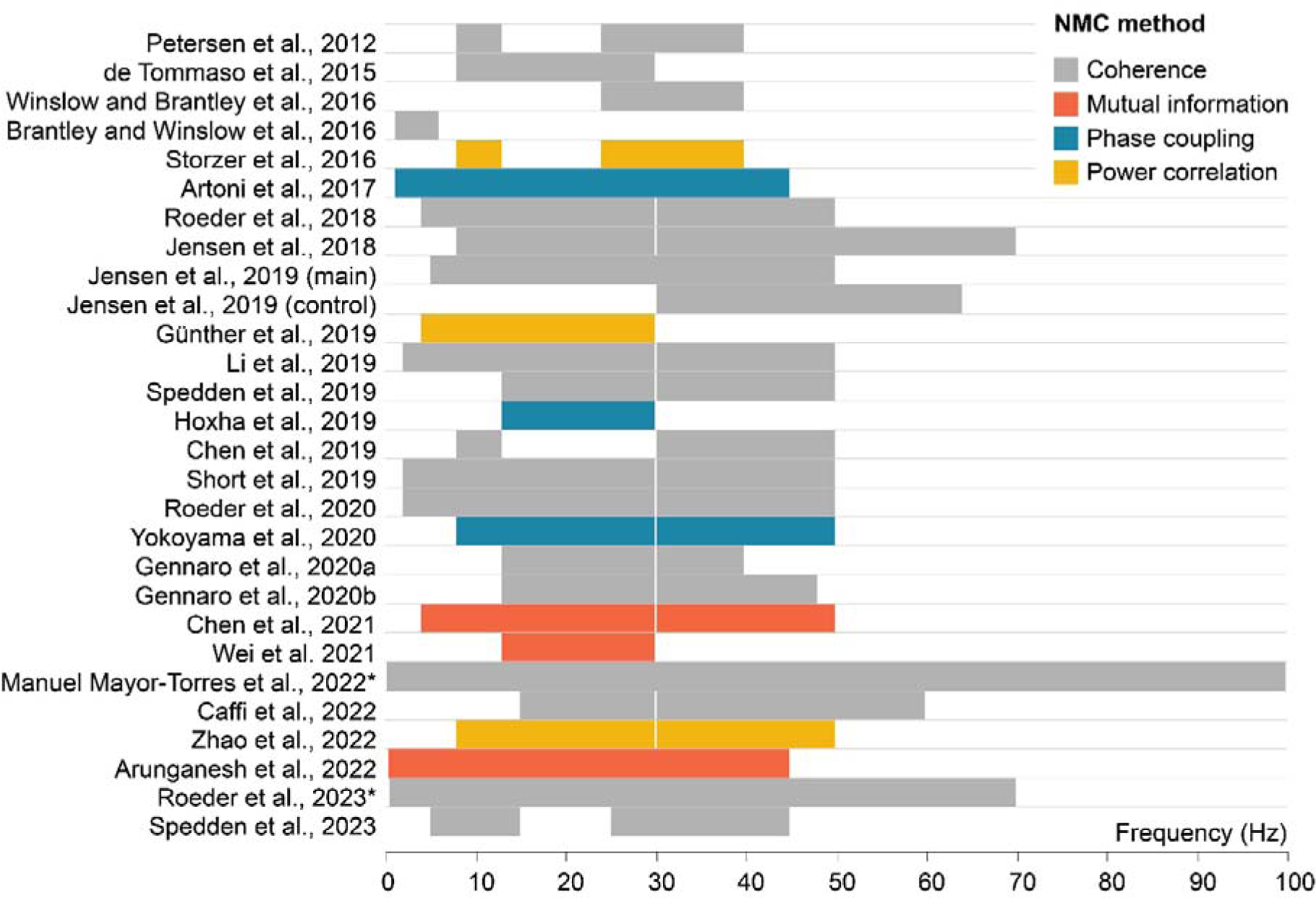
Method to quantify neuromuscular connectivity (NMC) and frequencies of interest per study. *These studies did not report specific frequency ranges of interest and therefore this figure is based on band-pass filter information.

Besides coherence, three studies opted to apply a power envelope correlation approach to quantify NMC. This means that a correlation was calculated between the power envelopes of the two signals (Hipp et al., 2012; O’Neill et al., 2015). Three other studies used a phase coupling method that quantifies whether the signals are time-locked (Lachaux et al., 1999). Lastly, three other studies used mutual information, which is a measure that assesses the correspondence in the value distributions of the signals. It is a useful method for examining non-linear relationships in the data, which may not be fully captured by coherence, power correlation, or phase coupling (Fraser and Swinney, 1986).

Most studies examined NMC in the beta [13-30 Hz] and gamma frequency [30-50 Hz] ranges. Thirteen studies examined the alpha frequency range [8-13 Hz], six studies the theta frequency range [4-8 Hz], and four studies the delta frequency range [1-6 Hz]. Three studies examined a broad frequency range but did not divide the data into different frequency bands. Three studies examined the 24-40Hz frequency range.

### 3.8. Neuromuscular connectivity modulations during the gait cycle

In general, the gait cycle is classified into eight gait phases (figure 8A). The first five phases comprise the stance phase. This stance phase starts with heel strike and ends with toe-off of the ipsilateral leg. The stance phase is further divided into two double support phases during which both feet are in contact with the ground (i.e. loading response and pre-swing) and two single support phases, during which only one foot is in contact with the ground (i.e. midstance and terminal stance). The stance phase encompasses approximately 60% of the gait cycle. The remaining 40% is allocated to the swing phase, further subdivided into initial swing, mid-swing, and terminal swing.

**Figure 8.**
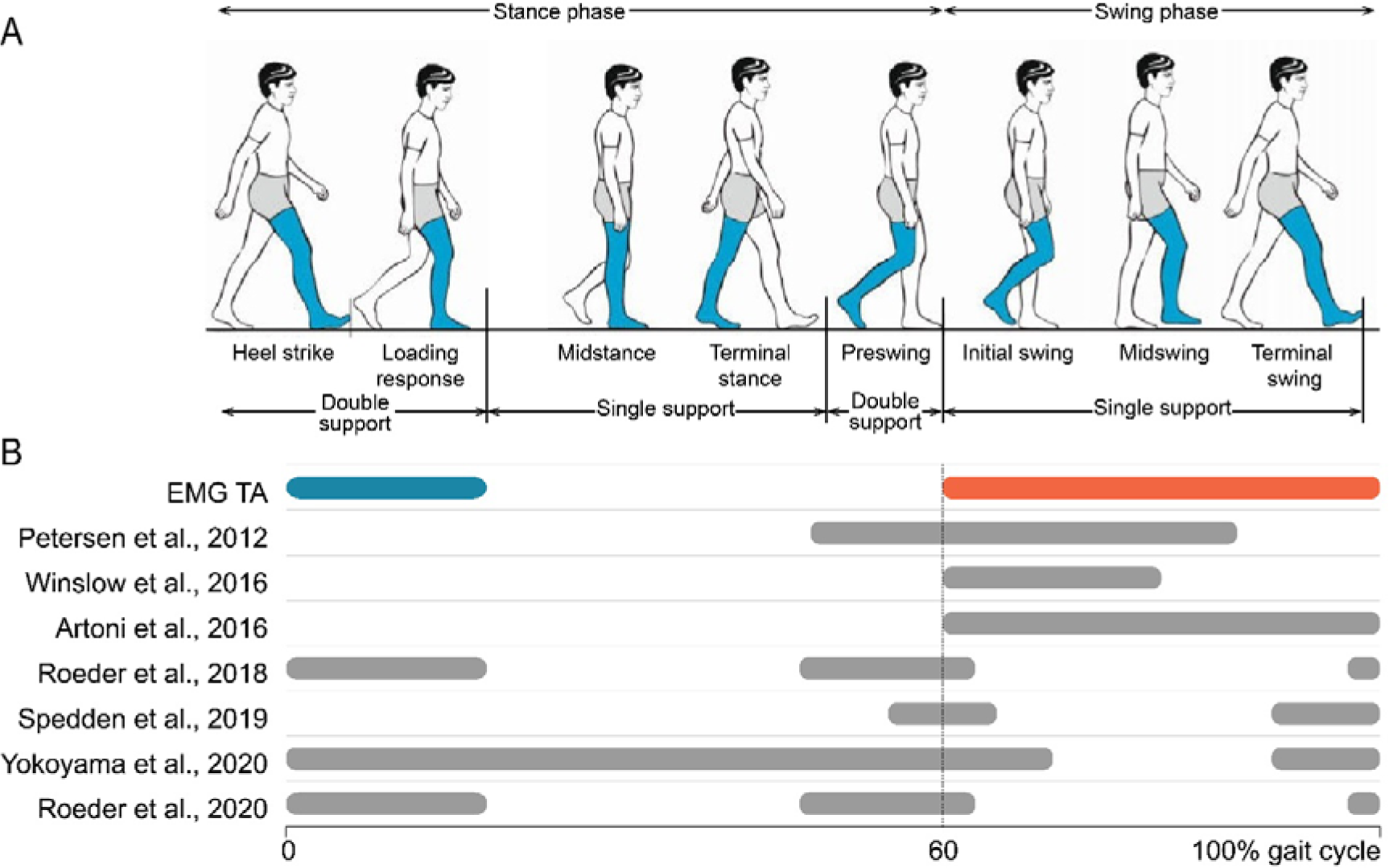
Neuromuscular connectivity (NMC) to tibialis anterior muscle (TA) over the gait cycle. A) Eight phases of the gait cycle focused on the right leg (image edited from Pirker and Katzenschlager, 2017). B) Activity of the tibialis anterior muscle over the gait cycle. The blue line represents eccentric activity, orange line represents concentric activity. Grey lines represent significant NMC.

#### 3.8.1. Tibialis anterior (TA)

The tibialis anterior muscle (TA) works eccentrically shortly after heel strike to control the toes’ descent. Its activation during the stance phase remains minimal. A secondary burst of activity emerges in the swing phase, requiring concentric action to lift the foot off the ground against gravity.

Figure 8 provides a comprehensive summary of studies investigating significant NMC to TA during various phases of gait. It includes results exclusively from studies involving healthy adult participants and those that distinctly differentiate between different gait phases. In cases where only the timing before heel strike was reported, the gait phase was estimated based on cadence and walking speed.

Multiple studies reveal prominent NMC in the swing phase of the gait cycle, primarily at beta and gamma frequencies ranges. The most substantial NMC was observed in the initial and terminal swing phases, which correspond to the periods following toe-off and preceding heel strike. These phases aligned with peak muscle activations.

However, Roeder et al. presented a contrasting view by reporting a lack of NMC during the swing phase (Roeder et al., 2020, 2018). Instead, their findings indicated NMC during the double support phases of gait. The first double support phase encompasses heel strike and loading response, coinciding with the eccentric action of the TA to control the downward movement of the foot. The second double support phase, pre-swing, involves foot preparation for toe-off. Apart from the studies of Roeder et al., some other investigations also identified significant NMC in these double support phases, with the largest values found in lower frequency bands.

#### 3.8.2. Gastrocnemius (GC) and soleus (SOL)

The main function of the soleus (SOL) during gait is the control of the anterior translation of the tibia in the stance phase. During this phase, the SOL works eccentrically. Conversely, the gastrocnemius muscle (GC) predominantly functions concentrically. It engages especially during the latter part of the stance phase, facilitating the propulsive advancement of the body. Jensen et al. found a strong NMC pattern in the gamma frequency range (approximately 40 Hz) between the brain region Cz and SOL, which persisted as long as SOL muscle activity remained active (Jensen et al., 2019). This distinct NMC faded as the stance phase ended. In parallel, a similar gamma range NMC was observed between Cz and GC during the stance phase, though this NMC was less pronounced compared to the Cz-SOL NMC. Instead, GC NMC emerged at both lower and higher frequencies around the time of push-off.

In the study of Yokoyama et al. significant NMC to GC was found mainly during the stance phase and extended into the initial swing (Yokoyama et al., 2020). In addition, Roeder et al. examined NMC to SOL and GC (Roeder et al., 2023). Similarly to TA, they found significant NMC during the double support phases of gait. While the presence of NMC aligned with GC activity during the pre-swing phase seems reasonable, coherence during the loading response phase raises questions, considering the lack of activation in both the GC and SOL during that period.

## 4. Discussion

The findings of this systematic review highlight the potential of neuromuscular connectivity (NMC) analysis in advancing our knowledge of cortical involvement in gait control. The vast majority of the included studies were able to identify significant NMC during walking. This reflects the presence of communication and interaction between the cortex and muscles (Conway et al., 1995; Mima and Hallett, 1999). However, the studies varied considerably in terms of methodologies and offered diverse interpretations of their findings. In this section, we will delve into these variations and their implications.

### 4.1. Neuromuscular connectivity modulations over the gait cycle

NMC is modulated over the course of a single gait cycle. It is hypothesized that these modulations align with muscle activation patterns. When considering the tibialis anterior (TA) for instance, significant NMC has been found during the swing phase as well as during the loading response (Artoni et al., 2017; Petersen et al., 2012; Roeder et al., 2020, 2018), two phases in which the TA is primarily activated (Bonnefoy-Mazure and Stéphane Armand, 2015). Similarly for the soleus muscle (SOL), significant coherence was found during the stance phase, while the gastrocnemius (GC) shows the largest NMC in the push-off phase of gait (Jensen et al., 2019). Some studies, however, fail to find NMC at the precise timing of muscle activation. For instance, Roeder et al. did not find significant NMC for the TA muscle during swing. In fact, they observed NMC to the TA, GC, and SOL muscles to be significant only for the double support phases. In addition, there was no difference between NMC to the ipsilateral and contralateral side of the brain (Roeder et al., 2023, 2020, 2018). These findings suggest that muscle activity alone does not fully explain NMC modulations and that there might be different mechanisms at play.

Communication over the corticospinal tract not only encompasses motor commands but also sensory information. This sensory information could be a contributing factor to NMC (Witham et al., 2011). This hypothesis has been tested by Riddle and Baker et al. (Riddle and Baker, 2005). After cooling down the arms of the subject, they found an additional time delay in the signal transmission between the brain and the muscles. The conduction time was roughly twice the normal conduction time in one direction. From this observation, it was concluded that both ascending (sensory) and descending (motor) pathways contribute to the occurrence of NMC. During gait, studies looking into the direction of information flow over the corticospinal tract found some cases during which the information flow was reversed, i.e. from muscle to brain. For instance, Petersen et al. found ascending information around 10 Hz (Petersen et al., 2012) and Jensen et al. reported that SOL activity preceded brain activity during the push-off phase of gait (Jensen et al., 2019). It is therefore possible that the increased NMC in the double support phase in the studies of Roeder et al. could be ascribed to increased sensory information as a result of ground contact (Roeder et al., 2023, 2020, 2018). Remarkably in the studies of Roeder et al., participants were walking barefoot. This could potentially lead to an increased sensory information flow during the stance phase.

While descending information has been linked to beta and gamma frequency ranges, this ascending sensory information has been related to lower frequency ranges, i.e. below 13Hz (Mehrkanoon et al., 2014; Mima et al., 2000). In the context of gait, we made similar observations. Instances where the EMG signal preceded the EEG signal were commonly linked to lower frequency ranges (Brantley et al., 2016; Petersen et al., 2012). In addition, in certain clinical populations experiencing difficulties with sensorimotor integration, a significantly lower NMC in the alpha band was found compared to age-matched controls (Almeida et al., 2005; Yokoyama et al., 2020). This supports the link between alpha NMC and sensory feedback. Other studies, however, hypothesized that these alpha waves originate at the spinal level. More specifically, studies conducted on patients following spinal cord lesions revealed intra- and intermuscular NMC in the alpha frequency band, with a notable reduction in the beta and gamma ranges (Barthélemy et al., 2010; Hansen et al., 2005; Norton and Gorassini, 2006). In addition, other studies posited that NMC in very low-frequency ranges (< 5 Hz) likely signifies the periodic modulation of the EMG envelope (Halliday et al., 2003). This perspective underscores that such synchronization may be more indicative of the overall patterns of muscle activity rather than distinct neural processes. In general, we can conclude that beta and gamma frequency ranges are linked with efferent input and alpha with afferent input. NMC in the delta and theta frequency ranges on the other hand may not signify true synchronization over the corticospinal tract.

### 4.2. Inconsistency in research design and methodology

The vast variety of methods used in the different studies makes it virtually impossible to pinpoint the exact causes of discrepancies between findings. Differences arise in the *gait task specifics*, including variations in surfaces, gait speed, and footwear conditions. Additionally, the inclusion or exclusion of specific gait phases varies strongly among studies. In fact, seven studies excluded the gait phases containing heel strikes from their analysis. This precaution is taken to avoid potential false NMC arising from the collision of the foot with the ground, introducing substantial artifacts in both EMG and EEG signals. Therefore, less information about NMC is available for the period surrounding the heel strike. Comparing gait-phase modulations is further complicated by the fact that some studies measure NMC in relation to the time before the heel strike. This approach lacks precision in specifying the exact gait phase that is being observed. The diversity in EEG setups, ranging from two to 128 electrodes, and disparities in EEG and EMG preprocessing steps add to the complexity.

Recent advancements in *EEG technology* have improved our ability to pinpoint the source of neural activity in the brain. EEG source localization can be used in walking studies, allowing for a more detailed understanding of the different brain regions involved in different phases of the gait cycle (Li et al., 2019). Nonetheless, most of the studies included in this review quantified NMC between EEG signals at the sensor level, being either the Cz electrode or a unilateral electrode over the sensorimotor cortex. Only four studies used the signal in the source space. Using source localization, Zhao et al. showed NMC not only between the muscle and the primary motor cortex, but also between the muscle and the premotor cortex, posterior parietal cortex, and cerebellum (Zhao et al., 2022). This approach holds promise for providing a more comprehensive understanding of the complex sensorimotor system by precisely identifying the EEG signal sources.

Different *methods of calculating NMC* have been adopted, each having its unique advantages and disadvantages. Coherence is the predominant method employed in the studies analyzed in this review. It is a frequency-based approach that assesses the similarity in the frequency characteristics between two signals. However, significant brain oscillations typically occur within a frequency range of 1-50Hz, while muscle activity ranges from 20 to 200Hz. As a result of these incongruent frequency ranges, using coherence may potentially lead to an underestimation of the NMC level. A similar consideration applies to phase coupling techniques, which examine whether a stable time delay can be found between EMG and EEG signals considered at the same frequency. To solve the problem associated with the use of coherence and phase coupling techniques, particular studies correlate the power envelope of the whole EMG signal with the power envelope of the EEG signal in specific frequency bands. This approach, however, requires the availability of long EMG/EEG recordings such that the correlation is calculated on enough independent data points. Coherence and power correlation are linear techniques, but studies showed that NMC in the sensorimotor system is non-linear (Yang et al., 2018). Therefore, the use of non-linear NMC approaches can be suggested, such as mutual information or cross-frequency coupling. Future research should therefore carefully consider different analysis techniques depending on the specific research question.

In summary, the diverse methods employed in various studies hinder direct comparisons of results. Nevertheless, the opportunity for valuable comparisons arises when specific gait tasks or population groups are investigated within the same study while maintaining consistent methodologies. Within this review, several studies undertook such comparative approaches, examining NMC across different population groups or walking conditions. These studies provided crucial insights for identifying overarching trends related to aging, pathologies, and walking conditions, which we will delve into in the following sections.

### 4.3. Altered NMC in aging and motor disorders

Examining NMC in an aging population can help us understand the significant impact of the aging process on the neuromuscular system. Aging is characterized by a reduction in motor unit numbers, modifications to the morphology and characteristics of existing motor units, and adjustments in inputs originating from peripheral, spinal, and supraspinal sources (Hunter et al., 2016). The impact of these degenerative processes on gait has been studied using NMC analysis.

The impact of age on NMC tends to vary depending on the type of motor task examined. Younger adults exhibit higher NMC in studies focusing on gross motor movements such as ankle contractions, elbow flexion contractions, and postural control perturbations (Bayram et al., 2015; Ozdemir et al., 2018; Spedden et al., 2018; Yoshida et al., 2017). Conversely, studies involving fine motor control tasks such as finger muscle contractions report higher NMC in older adults (Johnson and Shinohara, 2012; Kamp et al., 2013). When it comes to gait specifically, results also depend on the tasks examined. Studies involving treadmill walking demonstrate higher NMC in young adults, while this is not the case in overground conditions (Caffi et al., 2022; Gennaro and de Bruin, 2020; Roeder et al., 2020; Spedden et al., 2019; Yokoyama et al., 2020). The distinction in sensorimotor control during treadmill walking compared to overground gait could provide a potential explanation for these variations. These findings underscore the importance of considering the specific motor task under investigation when interpreting the results of NMC studies in the context of aging.

NMC analysis also demonstrated its ability to discriminate between a healthy population and patients with neurological or neuromuscular disease. Yokoyama et al. reported decreased NMC in PD patients compared to healthy controls, whereas Roeder et al. found no significant difference (Roeder et al., 2020; Yokoyama et al., 2020). Remarkably, in the study of Yokoyama et al. participants were tested in the “off-medication” state while in the study of Roeder et al. patients were optimally medicated. This relation with medication status was confirmed in other studies (Salenius et al., 2002). In stroke and MS patients, a reduced NMC during gait was found. This is in line with research showing reduced NMC during other motor tasks in these patients (Chen et al., 2018; Fang et al., 2009; Krauth et al., 2019; Mima et al., 2001). In CP patients, NMC patterns are different during gait compared to typically developing adolescents (Gennaro et al., 2020; Short et al., 2020). Higher NMC patterns are furthermore detected in sarcopenic older adults compared to age matched controls (Gennaro et al., 2020). These findings may potentially indicate a compensatory mechanism.

The hypothesis that patients are augmenting NMC as a compensatory mechanism implies the potential for modulation of NMC over a longer period. In stroke patients, NMC was observed to rise following a four-week turning-based treadmill training program (Chen et al., 2019). Interestingly, the larger the changes in NMC were, the better the recovery of gait symmetry. This is in line with previous research showing a relation between increments in NMC with increments in functional recovery (Zheng et al., 2018). Apart from modulation in the long term, particular studies also point towards an acute modulation of NMC in relation to task demands. Older adults exhibited an increase in NMC when faced with more demanding tasks, such as performing cognitive dual tasks or performing visually guided gait (Caffi et al., 2022; Spedden et al., 2019). This finding suggests that NMC tends to rise with increased engagement of cortical processes, particularly among older individuals. Overall, elderly and clinical populations tend to exhibit decreased NMC compared to a healthy, young population, but they are still capable of enhancing NMC when faced with higher task demands.

### 4.4. Task-dependent modulations of NMC

Apart from differences between populations, also the specific motor task demands should be considered. Significant NMC is typically found during sustained, isometric contractions, while NMC in the beta range is suppressed during dynamic motor tasks (Kilner et al., 2003; Omlor et al., 2007). Indeed, Petersen et al. compared NMC during static dorsiflexion contractions with NMC from Cz to TA during dynamic steady-state walking and confirmed this postulate (Petersen et al., 2012). Despite the dynamic nature of the task, NMC during gait was significant.

Besides the dynamicity of the task, it seems that NMC during gait is adjusted in response to the demands imposed on the sensorimotor network. Gait tasks that are more challenging for the sensorimotor network show a heightened level of NMC. This relationship has initially been found in studies involving cats (Armstrong, 1988; Drew et al., 1996; Drew and Marigold, 2015). When cats were required to navigate obstacles and carefully place their paws on ladder rungs, researchers observed a heightened firing rate in corticospinal neurons. Also in humans, an increase in NMC was observed during a visually guided walking task (Caffi et al., 2022; Jensen et al., 2018; Spedden et al., 2019). During this task, individuals must pay close attention to visual cues and make precise adjustments to their gait and foot placement based on visual input.

Likewise, slow walking showed higher NMC compared to fast walking (Chen et al., 2021; Wei et al., 2021). During slow walking, there is a conscious adjustment of step length, balance, and control of limb movements needed to ensure stability. Moreover, no significant NMC was observed in a study that exclusively focused on stepping movements without transitioning into continuous walking (Spedden et al., 2023). It was proposed that the heightened challenge of coordinating and initiating gait from a dynamic and inherently unstable position as seen during a steady-state gait, may impose additional demands on spinal and subcortical circuits.

In addition, two studies identified an increase in beta NMC in overground walking compared to treadmill walking (Roeder et al., 2020, 2018). According to the hypothesis, this would mean that overground walking places higher demands on the corticospinal network than treadmill walking. A decrease in cadence and increase in TA activity have been found during overground walking compared to treadmill walking (Lee and Hidler, 2008), suggesting that individuals indeed adapt their gait patterns in response to different walking surfaces. Additionally, in overground walking conditions participants were instructed to maintain a consistent, preferred walking speed. This requires them to consciously monitor their pace, hereby increasing the demand on the sensorimotor network. In contrast, it has been observed that treadmill walking necessitates enhanced dynamic stability and imposes higher demands on cortical neuromotor control (Herold et al., 2019; Yang and King, 2016). This clearly contradicts our initial hypothesis. It is worth noting that both studies comparing treadmill and overground walking did not exclude data from heel strikes. This raises the question whether artifacts related to heel strikes could potentially account for the observed NMC differences. Especially since these artifacts may be more pronounced during overground walking compared to treadmill walking (Riley et al., 2007).

Across these scenarios, it appears that variations in the task, which require greater engagement of the sensorimotor network, tend to be associated with elevated levels of NMC. However, increased beta coherence was observed during an isometric motor task requiring increased attention (Kristeva-Feige et al., 2002). This raises the possibility that the enhanced NMC in more complex tasks may be attributed to increased attention rather than larger sensorimotor network demands. To validate this hypothesis, additional research comparing varying degrees of sensorimotor demands and attention levels during gait is warranted.

### 4.5. Neuromuscular connectivity during gait: evidence for cortical control?

The interpretation of NMC results is constrained not only by variations in methodology, population groups, and gait task specifics but also by the fact that the actual physiological implications of NMC remain incompletely understood. The presence of NMC during gait is considered to provide evidence for the active role of the cortex in modulating gait control. This active role of the cortex has been confirmed by studies exploring the direction of information flow, as they consistently report EEG signals to precede EMG signals (Artoni et al., 2017; Petersen et al., 2012; Winslow et al., 2016; Yokoyama et al., 2020). In some cases, however, information flow in the opposite direction has been detected (Jensen et al., 2019; Petersen et al., 2012). Therefore, it is advisable to consider not only the descending corticospinal pathway but also ascending neural pathways. In addition, local spinal circuitries as well as other descending pathways should be considered when interpreting NMC measures (Glories and Duclay, 2023; Matsuya et al., 2017). This is necessary because any of these factors could exert a direct or indirect influence on the observed NMC measure. In other words, NMC is more than just the product of efferent cortical drive.

Roeder et al. propose even another interpretation of NMC (Roeder et al., 2018). As they find both EEG and EMG signals during double support to be phase-locked to the heel strike event, they suggest the heel strike to be an external event that affects both brain and muscle activity independently. Consequently, they question if the NMC they observed genuinely reflects true corticospinal synchronization. Although the time lag between EEG and EMG could be an argument, temporal precedence does not necessarily imply a causal influence between the signals (Kamiński et al., 2001). Additional research is therefore needed to better understand the actual physiological relevance of NMC.

## 5. Conclusions

In this review, the presence of corticospinal communication becomes evident, as most studies included in the analysis consistently report significant NMC during walking. Moreover, studies that compare groups or tasks using the same methodology show that NMC measurements can discern differences between groups or specific gait tasks. However, comparing NMC across studies is challenging due to the diverse techniques employed. Hence, future research should prioritize methodological alignment to facilitate comparisons across a broader range of gait tasks with varying sensorimotor demands, as well as across different population groups. As Zhao et al.’s findings revealed NMC to other brain regions, future investigations should expand their focus beyond central electrodes, possibly adopting source localization. Furthermore, previous research clearly showed the impact of muscle fatigue and attention on NMC (Ushiyama et al., 2011). It is therefore crucial to examine how cognitive factors, fatigue or a lack of sleep can alter NMC measurements during gait. These future steps will significantly enhance our comprehension of corticospinal control in human locomotion.

## Author Contributions

M.S., D.M. and T.T.d.B. made substantial contributions to the conception of the work, drafted and revised the manuscript. All authors have read and agreed to the published version of the manuscript.

## Funding

The work was supported by the Research Foundation Flanders (grant no. G0F76.16N to DM) and the KU Leuven Special Fund (grant no. 3M220016 to TDB).

## Acknowledgments

The authors would like to thank Benedicte Vanwanseele and Jessica Samogin for their insightful discussions.

## Conflicts of Interest

The authors declare that the research was conducted in the absence of any commercial or financial relationships that could be construed as potential conflict of interest.

## Notes

### Competing Interest Statement

The authors have declared no competing interest.

